# From Pixel to Wave: A Geometric Complementary Code for Hierarchical Pixel-Based Morphometry

**DOI:** 10.64898/2026.04.13.718311

**Authors:** William A. Marcil

**Author notes:** **Corresponding Author**: William A. Marcil, MD, 7101 Newport Avenue, Omaha, NE 68152.

## Abstract

This paper introduces a geometric complementary code (GCC) that bridges discrete digital pixel graphics with continuous analog wave mechanics in a geometric morphometrics framework. By oscillating four shaded cubic pixels in a Cartesian grid, an emergent pattern resembling a face-centered cubic (FCC) unit cell lattice appears. This pattern is modeled first in lattice space— incorporating polarities of the sagittal, transverse, and coronal planes traditionally applied to Cartesian space—and subsequently in Cartesian space as a topographic medium.

In the Cartesian model, topographic values divide into rise values, where the grid converges toward elevated features along the Y-axis, and run values, where it flares within terrain dips. This produces a surface grid that undulates like a propagating wave. Across both models, a polar continuum emerges, oscillating between crossed and uncrossed polarities at micro- and macro-grid scales when applied at the topographic tile level. Each tile oscillates to generate counter-oscillatory perspectives across the macro-grid, dynamically shifting between approaching-point and vanishing-point modes.

The GCC functions as a hierarchical pixel-based morphometry. It begins with pixel-scale analysis in a single ZX plane, advances to atomic-scale resolution across four ZX planes, and extends to topographic tile-scale across 16 ZX planes. This progression reveals a geometric expansion of FCC patterns within a nested 2×2 matrix processor. Grounded in a consistent Pythagorean grid-count relationship, the framework maps discrete pixel states onto continuous wave-like surface behavior.

By addressing limitations in current geometric morphometrics—where 3D scanning and semilandmark methods remain anchored in discrete landmarks or sparse points rather than detailed continuous topographic dimensions—the GCC offers a novel hierarchical bridge between digital and analog domains.

## Introduction

Geometric morphometrics (GM) is a powerful multivariate method for analyzing biological shape using Cartesian coordinates of landmark data. Over the past thirty years, Cartesian grids and associated statistical tools have become central to GM, enabling robust quantification of morphological variation through homologous landmarks and Procrustes superimposition (1,2).

Traditional GM approaches rely on discrete anatomical landmarks or semilandmarks to construct morphological datasets. While effective for many applications, these methods face notable limitations, including sparse sampling, challenges in establishing true homology, and difficulty fully capturing complex continuous surfaces (3,4). Recent integration of machine learning has enhanced pattern detection in morphological data, yet the geometric underpinnings of such algorithms often remain difficult to reverse-engineer (3).

Three-dimensional scanning technologies (structured light, CT, surface scanners, photogrammetry, and VR-assisted digitization) have extended GM toward surface-based studies (5-7). However, even these advances typically remain centered on discrete landmarks, outline curves, or sparse surface patches rather than detailed continuous topographic dimensions such as curvature, relief, rise/run dynamics, or wave-like undulations (8-10). Semilandmarks improve representation of curves and surfaces where homology is ambiguous, but they still emphasize geometric relationships over treating topographic features as independent continuous variables (8,11). Emerging surface-based and landmark-free methods offer richer detail, yet no existing framework provides a hierarchical pixel-based morphometry that explicitly maps discrete Cartesian pixel states to continuous analog wave behavior via an emergent FCC lattice and polar continuum.

The present work introduces the geometric complementary code (GCC) to address this gap. By oscillating four shaded cubic pixels in a Cartesian grid with analysis performed on successive ZX planes, an emergent FCC-like lattice arises. This pattern is modeled first in lattice space using sagittal, transverse, and coronal polarities, then in Cartesian space as a topographic medium characterized by rise (convergence toward elevated Y-features) and run (flaring within dips) values. The resulting polar continuum oscillates between crossed and uncrossed states across micro- and macro-grids at the topographic tile scale, producing counter-oscillatory perspectives that shift between approaching-point and vanishing-point modes. Scaling hierarchically from single-plane pixel analysis through atomic (four-plane) and tile (16-plane) levels within a 2×2 matrix processor, the GCC is grounded in a Pythagorean grid-count relationship. This provides an explicit geometric construction for bridging discrete digital states with continuous topographic wave mechanics.

## Methods

The Geometric Complementary Code (GCC) implements pixel-based morphometry through a nested hierarchy of complementary pixel pairs. Analyses are carried out on successive ZX-planes in the adopted coordinate system, with the Y-axis representing the primary direction of topographic rise and run. This relationship is examined only after addressing the simpler pixel- and atomic-scale behavior.

At the pixel scale, four shaded pixels form a 2×2 matrix. This 2×2 matrix constitutes one quadrant that contributes to the emergent face-centered cubic (FCC) unit cell pattern, with the overall lattice encoded through variations in pixel shading within and across ZX planes (Figs. 1, 2). In the FCC lattice, six key positions align with the polarities of the sagittal, transverse, and coronal planes, providing a consistent set of opposing axes in Cartesian space (Figs. 1, 2).

**Figure 1.**
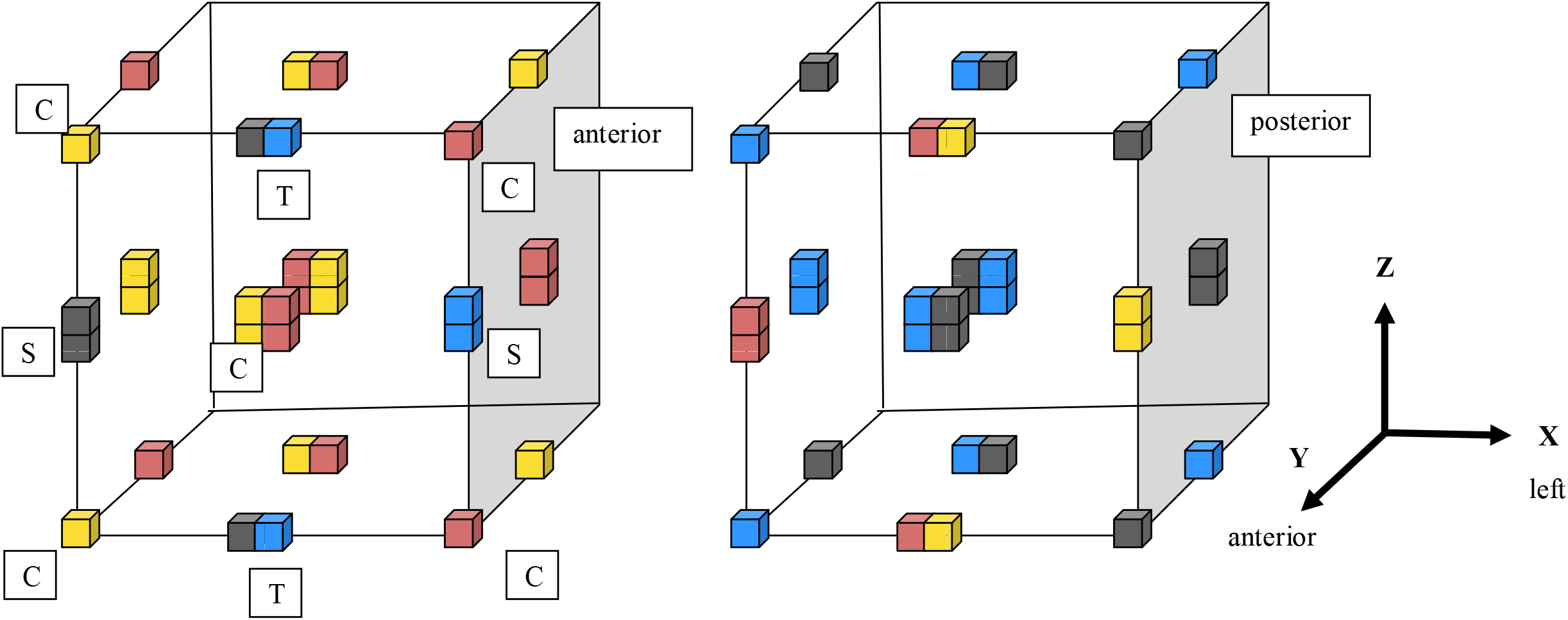
Lattice-space model across four successive ZX-planes (1–4, progressing anterior to posterior). Each ZX-plane contains a lattice unit cell composed of four quadrants, with each quadrant formed by a 2×2 arrangement of shaded pixels. Warm (gold, red) and cool (blue, black) shades encode complementary FCC-like sublattices analogous to sodium- and chloride-positioned elements. These sublattices align along the Y-axis, producing overlapping relational structures most clearly visible in ZX-planes 1 and 3. Dissection-plane polarities—coronal (center vs. corner, representing inside–outside polarity) and planar-side (sagittal vs. transverse)—govern the organization of each quadrant. Together, these complementary polarities generate the hierarchical relationships that give rise to the emergent face-centered cubic (FCC) pattern across the stacked ZX-planes, establishing the foundational relational unit for the Geometric Complementary Code.

**Figure 2.**
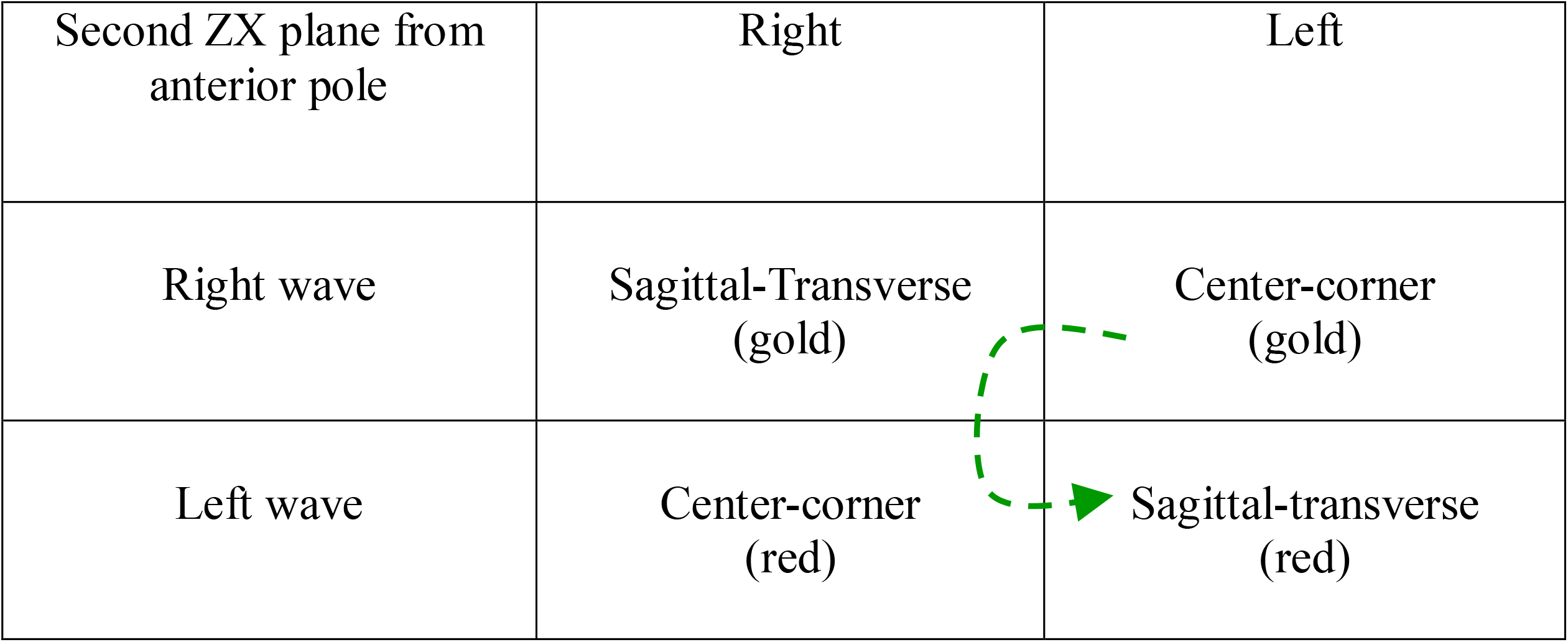
Pixel-scale 2×2 matrix (single ZX plane) illustrating the first hierarchical step of polar organization. The matrix shows the polar hierarchy of left and right waves, each composed of one coronal aspect and one planar-side aspect (transverse and sagittal), establishing the foundational complementary relationships within the GCC.

Dissection planes serve as standardized anatomical references: the sagittal plane divides left from right, the transverse plane divides superior from inferior, and the coronal plane divides anterior from posterior. When these polarities are applied to the FCC lattice via a 2×2 matrix, complementary relationships emerge naturally (Fig. 2). In this pixel-scale 2×2 matrix, the four cells function as poles defined by surrounding parameters. Coronal polarity pairs center and corner pixels, while planar-side polarity pairs sagittal and transverse elements within each quadrant (Fig. 1). This establishes equal and opposite poles: one pair defined by coronal (center– corner) relationships and the other by planar sides (sagittal–transverse) (Fig. 2). In coronal pairs, the corner serves as an external pole relative to the center, a property not present in sagittal or transverse orientations. This asymmetry enables a transformational step from a standard Cartesian grid to a diagonal lattice within three-dimensional space.

The hierarchy extends by synchronizing inverted right and left waves, allowing complementary opposites to form a polar continuum centered at the origin of the unit cell (Figs. 1, 3). Unlike the component waves arranged side by side, the polar continuum consists of two unified waves: one formed by phases that cross the vertical midline (“crossed”) and one formed by phases that do not cross (“uncrossed”) (Figs. 1, 3). These crossed and uncrossed states represent opposite ends of the polar continuum and cycle along the Y-axis.

**Figure 3.**
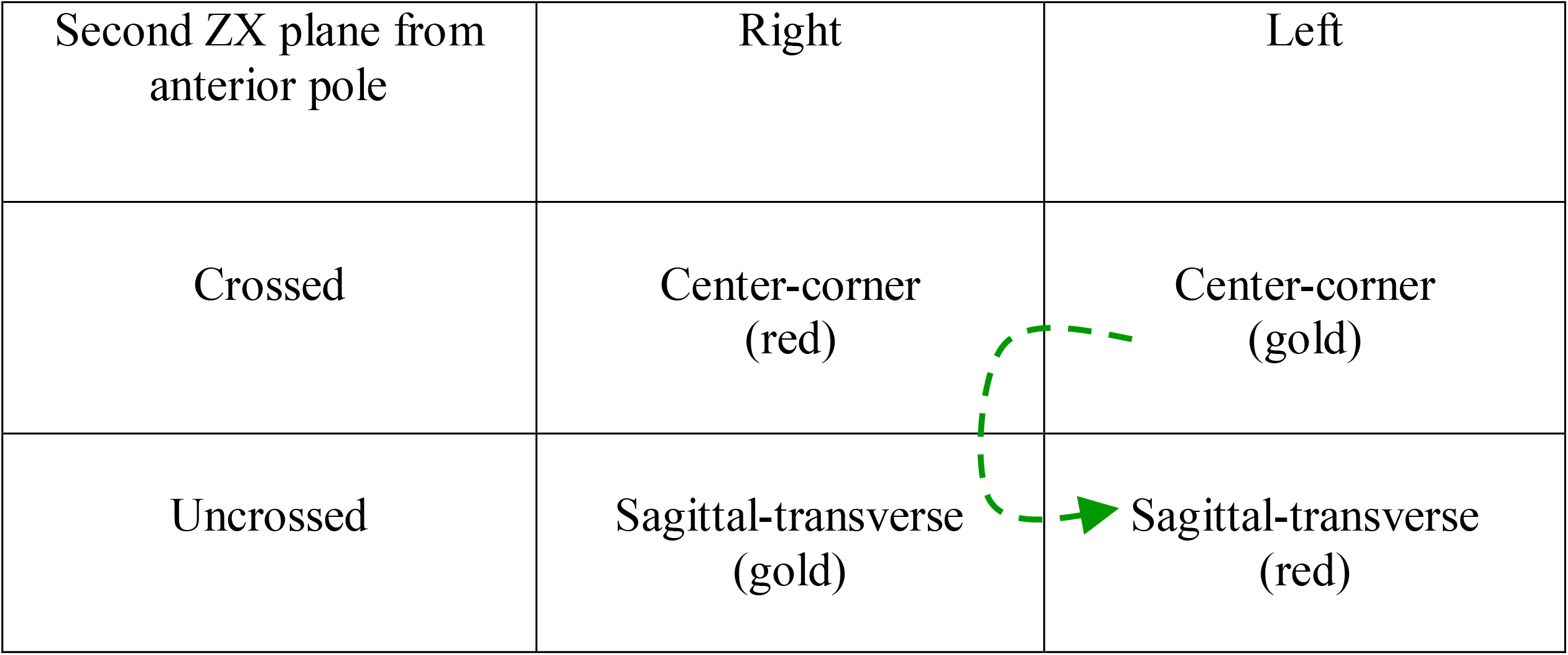
Pixel-scale 2×2 matrix (single ZX-plane) illustrating the second hierarchical step of the GCC. At this stage, coronal and planar-side polarities merge into a unified polar continuum centered at the unit-cell origin. This enmeshed configuration generates the first crossed and uncrossed wave components, which integrate the complementary pairs established in the initial hierarchy. The resulting crossed/uncrossed polarity forms the foundational relational structure from which higher-level wave behaviors emerge.

At the atomic scale, the model incorporates four ZX planes featuring distinct warm (Na^?^-like) and cool (Cl^?^-like) sublattices (Fig. 1). Warm pixels (red and gold) represent sodium atomic positions, while cool pixels (blue and black) encode chloride positions. For either warm or cool shades, the coronal center position forms a cubic arrangement of 8 pixels, surrounded by four square planar sides (each 4 pixels per side) and corner positions extending 2 pixels along the Y-axis (Fig. 1). In the 2×2 atomic matrix, warm pixels first merge along the Y-axis and then laterally as left and right waves (Fig. 4). Cool pixels follow the same pattern within their sublattice, resulting in two parallel polar continua positioned side by side along the Y-axis (Figs. 1, 4, 5).

**Figure 4.**
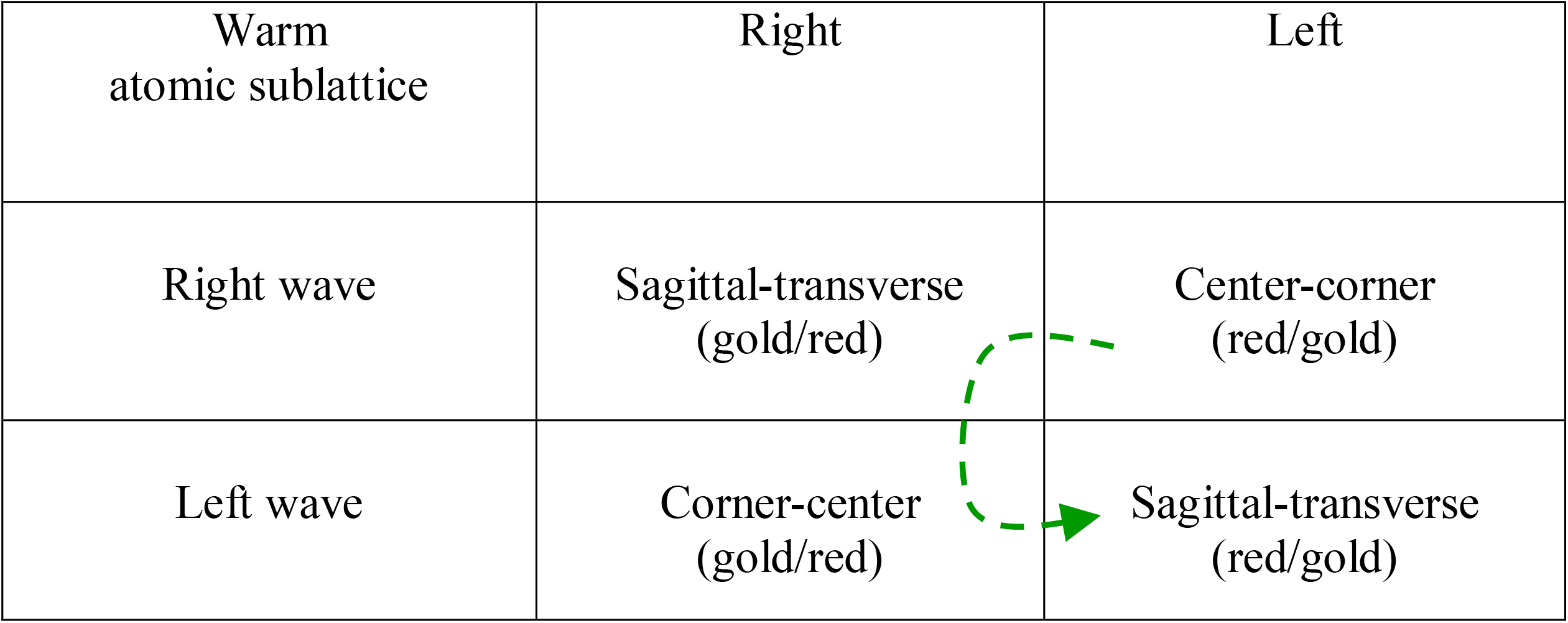
Atomic-scale 2×2 matrix spanning four successive ZX-planes, illustrating the first hierarchical step of relational polarity at the atomic level. Warm (gold, red) and cool (blue, black) pixels aggregate along the Y-axis to form complementary FCC-like sublattices analogous to sodium- and chloride-positioned elements. Within each sublattice, the coronal center pole expands into a cubic arrangement of eight pixels, each planar-side region contains four pixels, and the coronal corner positions extend two pixels along the Y-axis. These relational groupings establish the atomic-scale crossed and uncrossed configurations that parallel the pixel-scale hierarchy, providing the structural basis for higher-level wave organization within the Geometric Complementary Code.

**Figure 5.**
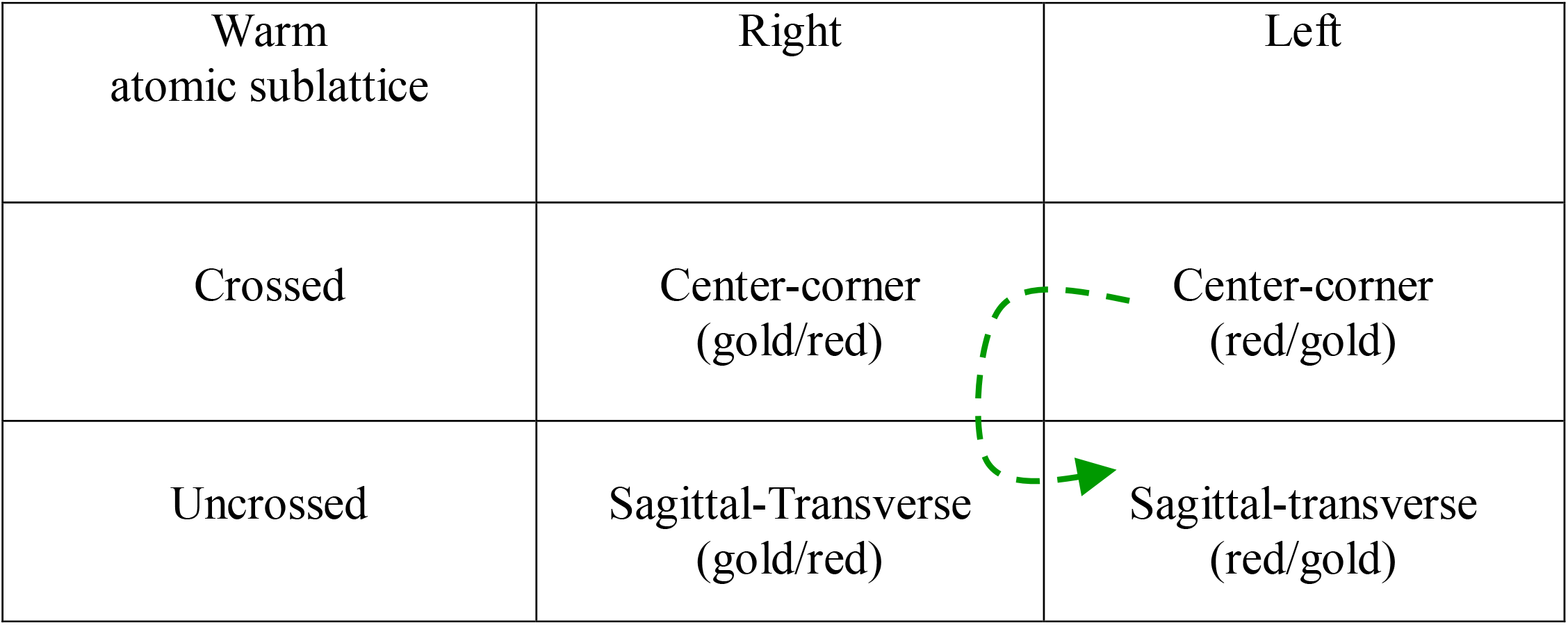
Atomic-scale 2×2 matrix spanning four ZX-planes, illustrating the second hierarchical step of relational polarity within the GCC. Within each warm and cool sublattice, pixel shades aggregate into right- and left-oriented waves that link coronal and planar-side polarities, forming an atomic-scale polar continuum. The two sublattices occupy anterior and posterior positions, extending the relational structure established at the pixel scale. This configuration demonstrates how the 2×2 matrix concept generalizes from a single ZX-plane (pixel scale) to four ZX-planes (atomic scale), applying the same complementary polar organization consistently across both two- and three-dimensional representations.

Generating left and right waves through coronal and planar-side polarity produces a unified polar continuum wave with crossed and uncrossed components centered at the unit cell origin (Figs. 4, 5). The atomic-scale hierarchy mirrors the pixel-scale hierarchy because transformations occur primarily along the Y-axis, while lateral unit-cell relationships remain unchanged. Consequently, Y-axis columns are scale-independent and provide the pixel contributions needed for expansion to higher organizational levels. At both scales, the polar continuum of crossed and uncrossed poles forms a system of mutually nested polarities that participate continuously in the diagonal grid (Figs. 3, 5).

To preserve left–right polarity throughout the hierarchy, a yin–yang pairing serves as the final common pathway for unification. In this context, yin–yang represents a dynamic balance between crossed and uncrossed waves, maintaining left–right distinction within the polar continuum while allowing complementary opposites to form a higher-dimensional whole.

## Results

### Encoding Topographic Dimensions

Topographic dimensions are quantified by analyzing pixel arrangements within triangular surface areas (Fig. 6). A sloped triangular surface preserves the incline of a 2D triangle (base to apex) while also exhibiting a 3D slope in Cartesian space. Such surfaces can be effectively represented in a two-dimensional medium due to their topographic properties (Fig. 7).

**Figure 6.**
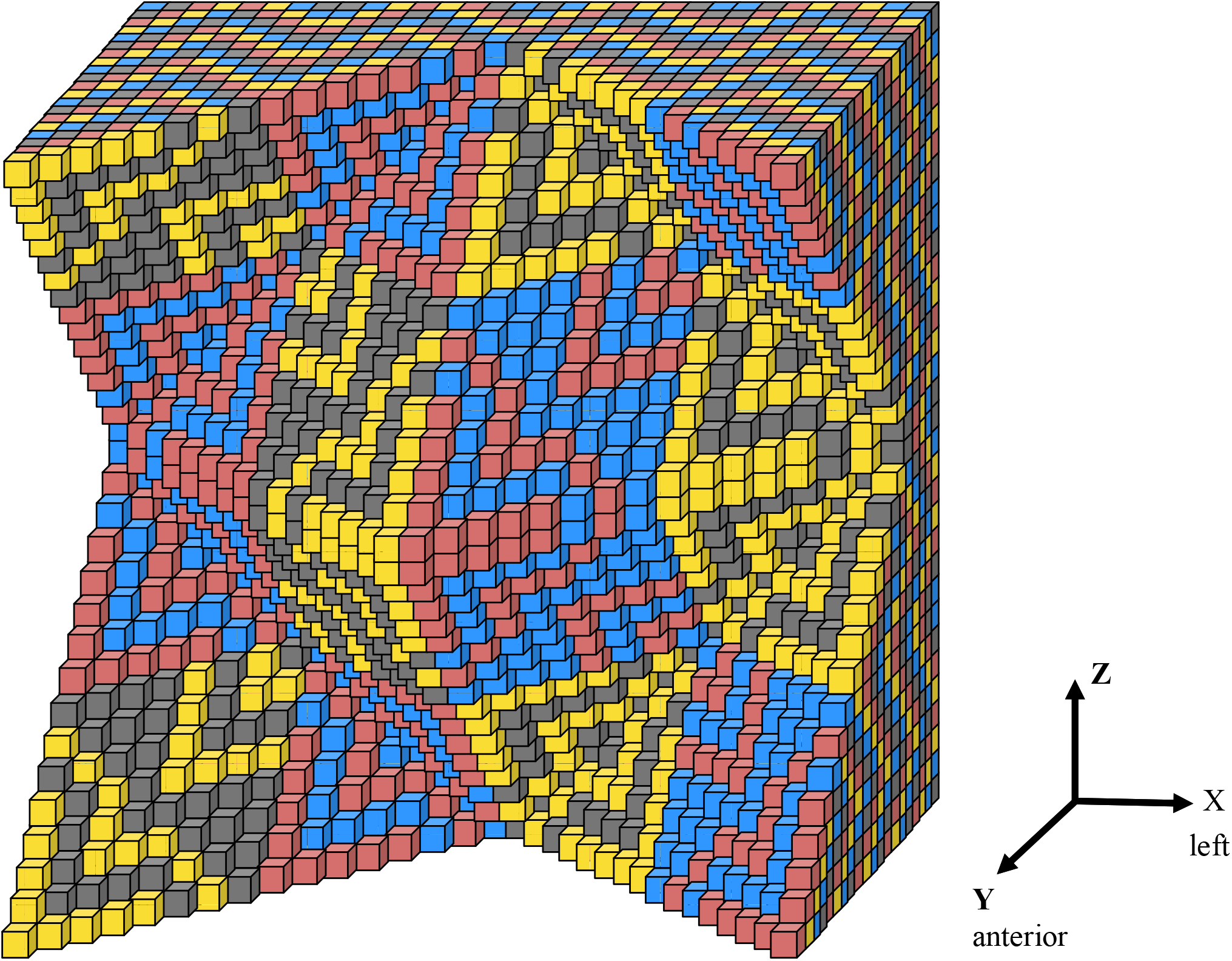
Topographic tile illustrating the relational organization of rise and run components. The central coronal pole (rise) protrudes along the Y-axis and is composed of four hexidecants, which transition into four planar-side regions (run) that retract into edge-like troughs (two hexidecants each). These planar-side regions connect to four corner positions that again protrude along the Y-axis (one hexidecant each). Color coding distinguishes the left-oriented wave (black and gold pixels), which displays planar-side polarity on the left and coronal polarity on the right, from the right-oriented wave (blue and red pixels), which exhibits the opposite arrangement. Synchronization of these left- and right-oriented waves produces a yin–yang distribution of reciprocal color coding between rise (coronal) and run (planar-side) poles, revealing the complementary structure that underlies the topographic hierarchy.

**Figure 7.**
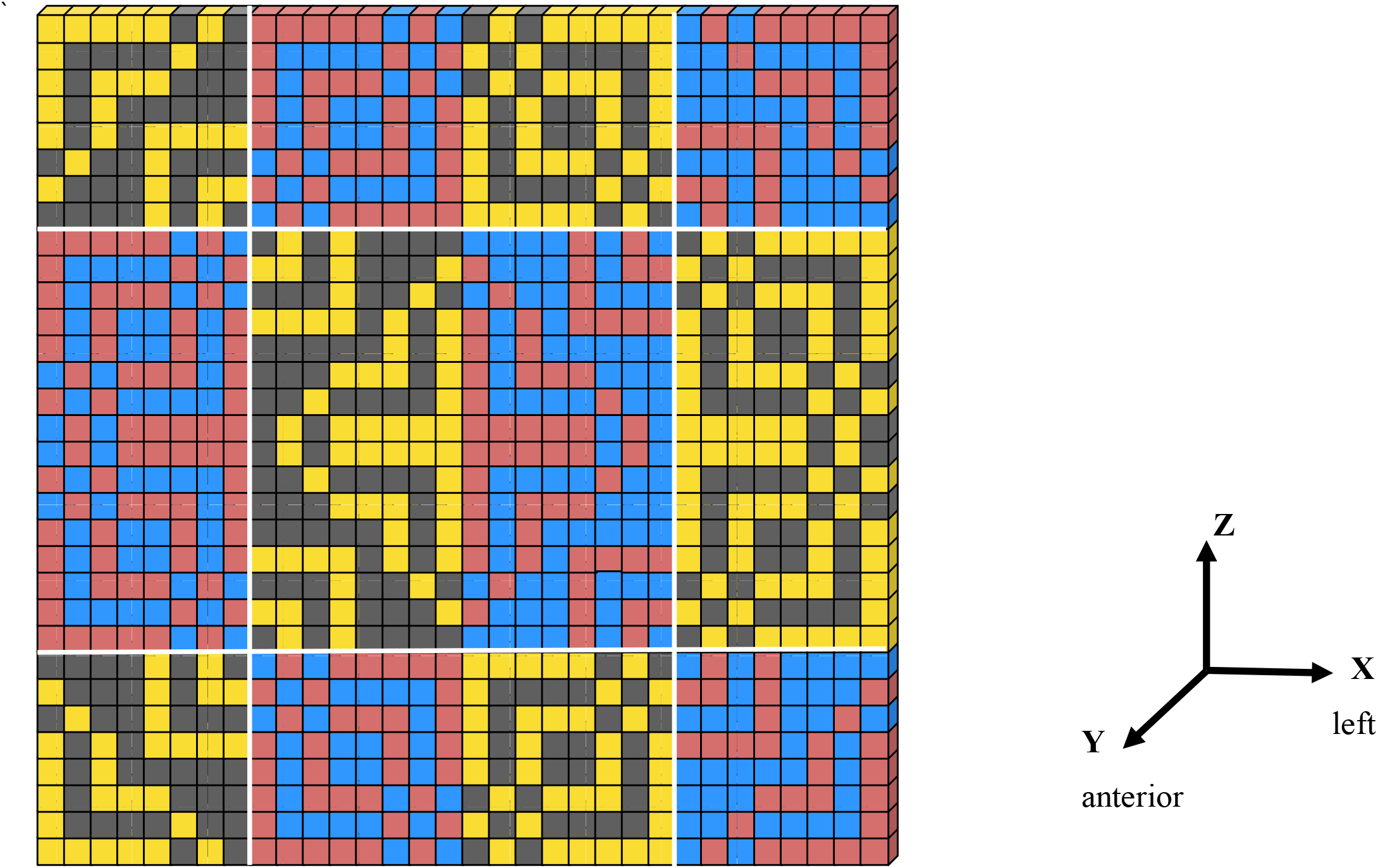
Two-dimensional representation of the topographic tile illustrating the polar continuum generated by hierarchical blending of right- and left-oriented waves. Alternating warm (gold, red) and cool (blue, black) shades radiate circumferentially from the center, encoding the relational transitions that occur as crossed and uncrossed phases nest across scales. This pattern reflects the pixel-level hierarchy in which right and left waves merge into a continuous topographic waveform, with internal right–left shading shifts distinguishing micro- and macro-scale continua of crossed and uncrossed polarities. The resulting distribution reveals how complementary opposites integrate into a unified relational structure within the GCC.

A topographic tile exhibits a polarity breakdown along its dissection planes that parallels the organization of an FCC unit cell, sharing the same landmark orientations. The topographic unit cell is divided into four quadrants, each further subdivided into four sections (hexidecants), analogous to pixels in a 2×2 grid (Figs. 1, 6, 7).

### Rise and Run Components

A rise aspect tapers to a point at opposite corners of the hexidecant, consistent with vertex morphology (Fig. 6). In contrast, the run feature appears as a diagonal flare bisecting the hexidecants, resembling an edge or base. A single quadrant can generate two adjacent, base-to-base triangles through the synchronized pairing of rise and run as complementary opposites (Fig. 6).

In this framework, rise and run values function as topographic XYZ measurements analogous to elevation data in standard mapping. A rise occurs when the value along the Y-axis increases more than the values in the perpendicular plane, producing a peak. A run occurs when the perpendicular plane values increase more than the Y-axis value, forming a trough. Rise and run thus serve as perpendicular components of a diagonal grid in 3D space.

### Hierarchical Wave Patterns and Polar Continuum

The 2×2 matrix dimensional hierarchy begins with opposing left and right waves, each consisting of one rise and one run phase (Figs. 7, 8). Synchronization of these waves generates a continuous pattern of nested crossed and uncrossed waves along the Y-axis, forming a coherent polar continuum (Figs. 7, 9).

**Figure 8.**
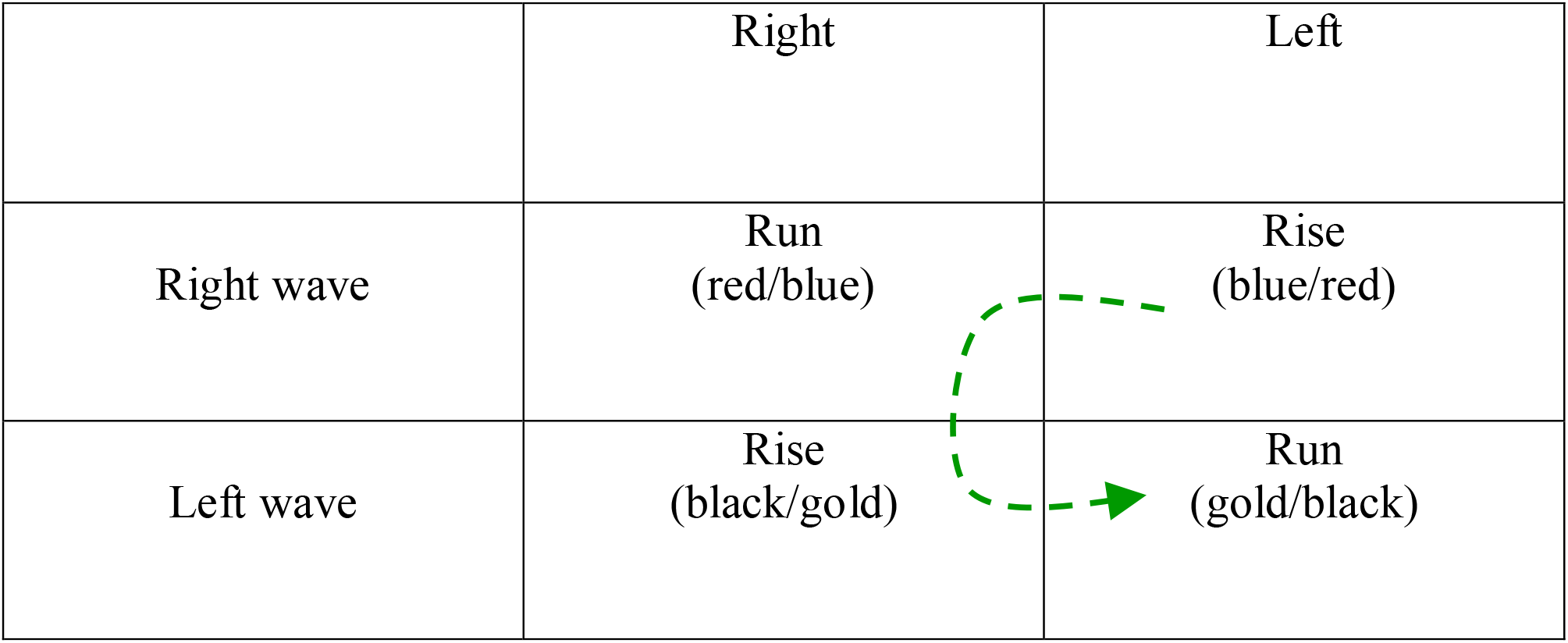
Tile-scale 2×2 matrix illustrating the first hierarchical step applied to the topographic tile. Following the same complementary polar organization established at the pixel and atomic scales, each tile contains opposing left- and right-oriented waves composed of one rise (coronal) phase and one run (planar-side) phase. This configuration demonstrates how the FCC-based relational hierarchy extends to the topographic scale, with the 2×2 matrix serving as the generative engine that propagates complementary opposites across increasingly complex surface structures.

**Figure 9.**
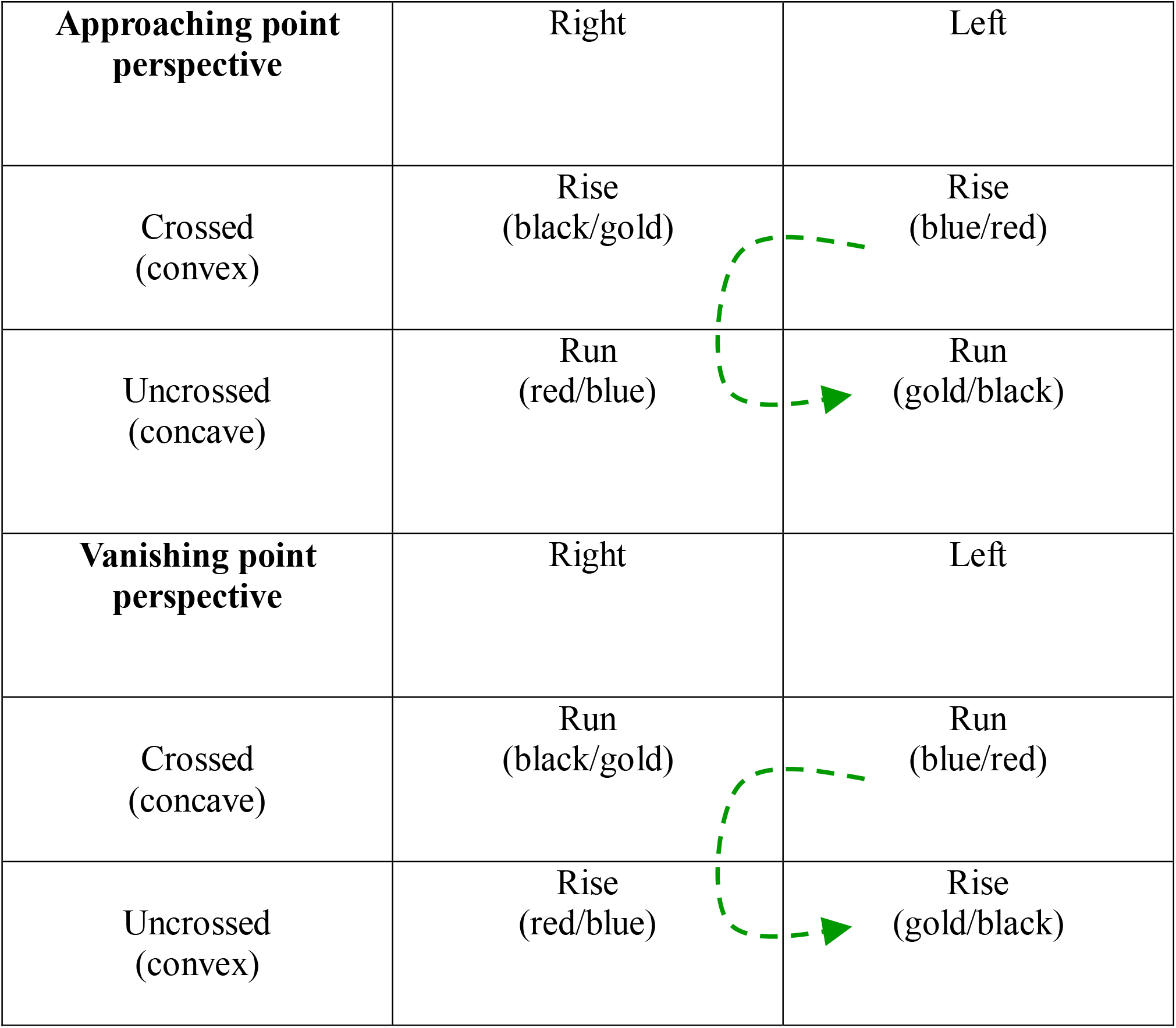
Tile-scale 2×2 matrix illustrating the second hierarchical step at the topographic tile level. Coronal (rise) and planar-side (run) polarities merge to form a polar continuum of crossed and uncrossed waves centered at the unit cell origin. The configuration exhibits a yin-yang pattern of reciprocal nesting, mirroring the hierarchical progression seen in pixel- and atomic-scale matrices but applied to the topographic surface and highlighting the self-similar nature of the GCC across scales.

Alternating warm and cool surfaces from the interior to the exterior of the tile illustrate the interplay between microscopic nesting and the larger FCC grid pattern (Fig. 7). For example, a warm uncrossed peak alternates with a cool crossed phase within the macro crossed wave.

Similarly, the macro uncrossed wave alternates between warm/uncrossed and cool/crossed phases, with right–left polarity reversed within each warm and cool division (Fig. 7). These mutually nested crossed and uncrossed phases create nesting at both micro- and macro-scales and produce a macro-level yin–yang pattern along the Y-axis, allowing each scale to influence the other (12) (Figs. 7, 9).

Rise-and-run polarities generate right- and left-oriented waves, which invert at a secondary polarity level. This produces a continuous polarity spectrum in which crossed and uncrossed states nest at both macro and micro scales, unifying into a macro yin–yang polarity. This hierarchical pairing of opposites—each level giving rise to the next—illustrates how opposites function as complementary pairs (13).

### Perspective Shifts and Surface Curvature

The topographic tile can oscillate between an approaching-point perspective and a vanishing-point perspective, depending on whether the rise and run polarities localize to opposite crossed or uncrossed macro positions (Figs. 6, 9, 10). In this model, rise phases correspond to convex surfaces, while run phases correspond to concave surfaces. Because convexity and concavity are fundamental concepts in geometric morphometrics, the GCC provides a valuable framework for their analysis (14). As with approaching point-perspective, a 2-D vanishing-point construction is possible because of the topographic dimension of the medium (Figs. 9, 11).

**Figure 10.**
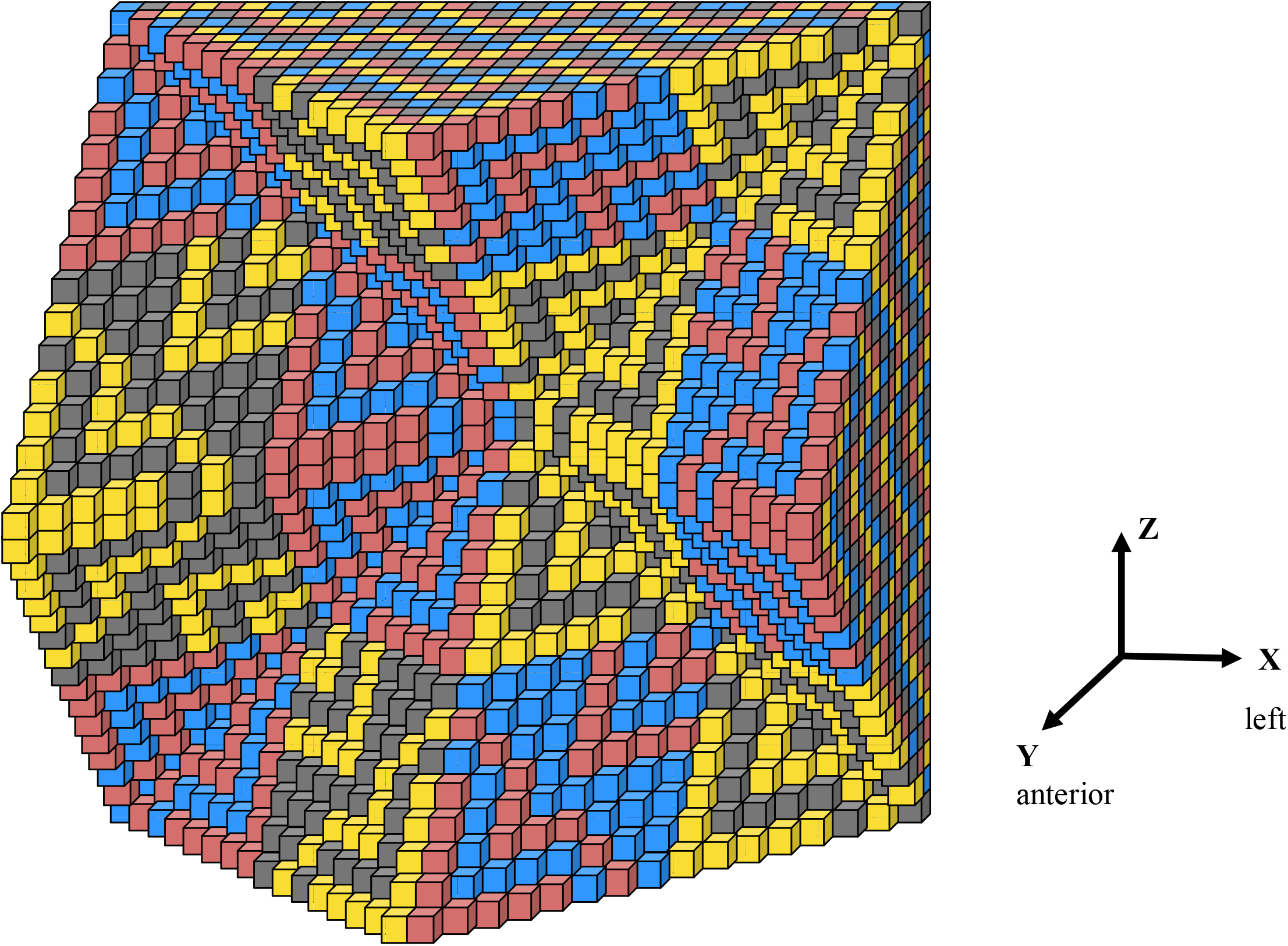
Oscillation of the topographic tile (see Fig. 6) illustrating how rise/run and crossed/uncrossed polarities interact across micro- and macro-scales. As rise (coronal) and run (planar-side) components exchange positions within the crossed and uncrossed macro-waves, the tile undergoes dynamic shifts between approaching-point and vanishing-point perspectives. This oscillatory behavior reveals how complementary phases reorganize within the polar continuum, producing hierarchical perspective changes that propagate through the GCC’s self-similar structure.

**Figure 11.**
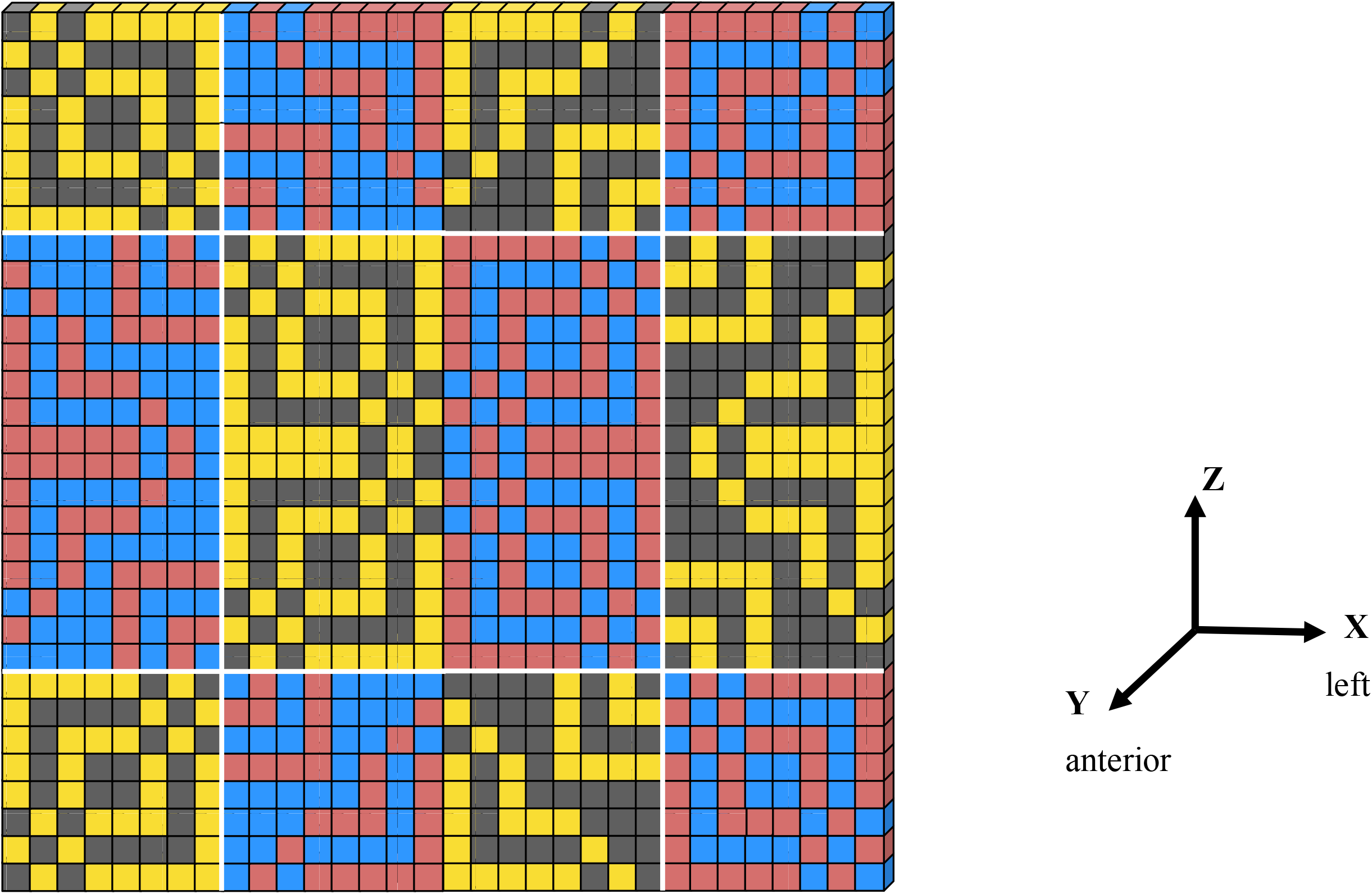
Two-dimensional representation of the oscillation shown in Figure 10, depicting the topographic tile’s dynamic shifts between rise/run phases and crossed/uncrossed polarities at micro- and macro-scales in a flattened view.

### Constructive Interference and Hierarchical Scaling

The juxtaposition of inverted right and left waves generates a constructive interference pattern within the tile structure. When two waves align in phase, their peaks and troughs add, producing constructive interference (15). In the same way, when right- and left-oriented topographic waves are in phase, their combined effects create elevated peaks in rising regions and deeper depressions in lowering regions (Figs. 6, 10). This interference forms a continuous pattern of nested crossed and uncrossed surface structures—analogous to pixel-scale models, but more clearly showing their surface polarization about the Y-axis and the resulting blurring of dimensional and polarity boundaries.

The FCC grid employs a 2×2 matrix to organize hierarchical relationships and generate continuous scaling. As the hierarchy of oscillating pixelations unfolds, each output becomes the 2×2 matrix input for the next level. Pixel-scale analysis requires a single ZX plane, atomic-scale analysis requires four ZX planes, and tile-scale analysis requires sixteen ZX planes (Figs. 6, 10). This produces geometric expansion by a factor of four at each step. The process culminates in two waves rotating around the Y-axis in a yin-yang pattern that alternates between crossed and uncrossed positions, generating a self-similar fractal-like structure.

## Discussion

The Pythagorean theorem, commonly expressed as *a*^*2*^ *+ b*^*2*^ *= c*^*2*^, finds a simplified geometric representation in the pixel-based medium of the GCC. Where pixels form square grid units, the relation becomes P + Q = R, with P and Q representing pixel counts along the legs of a right triangle and R the pixel count along the hypotenuse. This provides a direct geometric proof within the grid. In a topographic tile, P corresponds to the right-wave rise phase, Q to the right-wave run phase, and R to the left-wave structure (Fig. 12).

**Figure 12.**
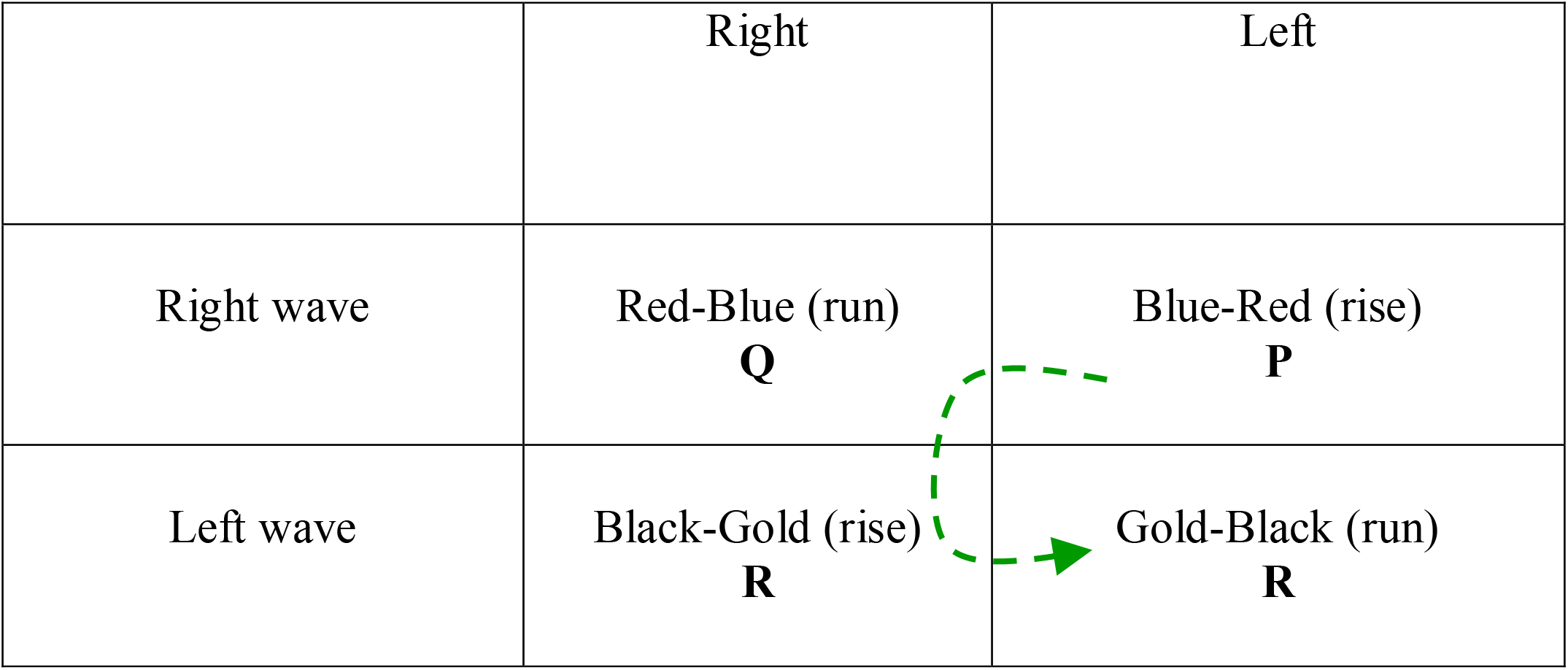
Geometric demonstration of the Pythagorean relationship within the GCC at the topographic-tile scale. The expression *P + Q = R* is shown in relational form, where *P* corresponds to the right-wave rise phase in quadrant I, *Q* corresponds to the right-wave run phase in quadrant II, and *R* represents the left-wave structure spanning quadrants III and IV. This configuration illustrates how the Pythagorean relationship functions as a hierarchical cycle within the 2×2 matrix, linking pixel-level geometry to topographic wave behavior and revealing the diagonal-grid logic that underlies the GCC’s self-similar structure.

Geometric dissection-plane polarities link the GCC to the Cartesian coordinate system, enabling complementary pairs to couple through the dimensional hierarchy. A geometric progression with factor 4 produces self-similar patterns across scales—from pixel to atomic to topographic— through a 2×2 matrix hierarchy applied recursively. This hierarchy progresses through successive polarities (perpendicular plane, Y-axis, crossed, uncrossed, and yin–yang). At the foundation of this hierarchy, dissection-plane polarities require reinterpretation to better account for surface areas and to establish the diagonal grid geometry governed by the Pythagorean relationship.

The surface possesses its own hierarchical capacity and can be considered quasi-independent of the underlying grid. When an approaching-point topographic tile is nested into an extended tile composed of four additional approaching-point tiles, a recursive hierarchy of oscillations emerges across all surface scales (Fig. 13). This hierarchy consists of four approaching-point tiles bisected by a vertical white line, followed by two vanishing-point tiles arranged as columns and bisected by horizontal white lines. The sequence culminates in a single approaching-point tile at the center of the extended tile—an oscillating derivative of the originals, notably not bisected by a white line (Fig. 13). This shows that the 2×2 matrix engine operates within the surface itself, exhibiting wave-like continuity rather than the discrete, particle-like behavior of individual pixels.

**Figure 13.**
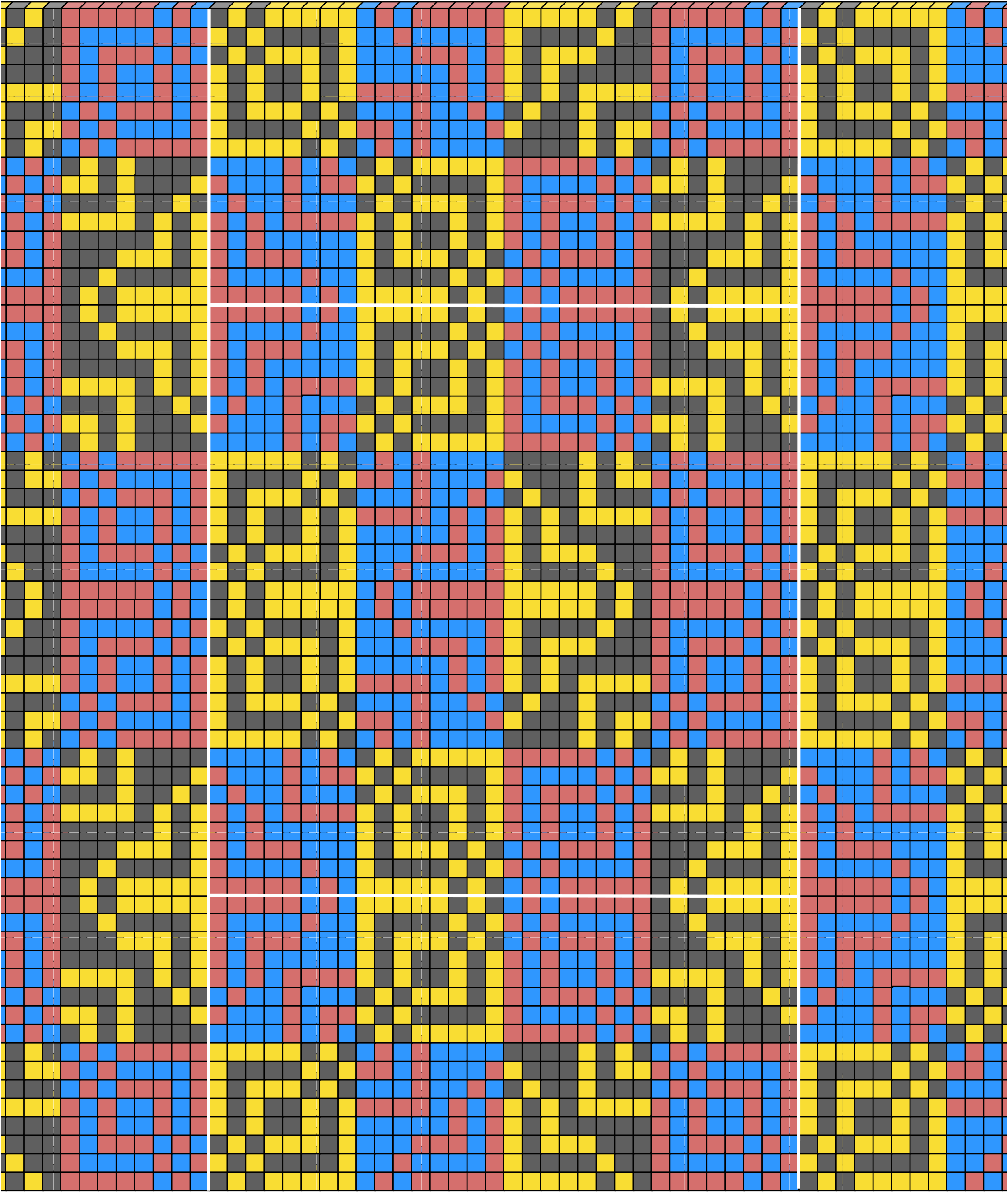
Oscillatory hierarchy of the topographic-tile surface illustrating recursive nesting generated by the GCC’s 2×2 matrix engine. Four approaching-point tiles, each bisected by a vertical white line, form the outer layer of the hierarchy. These are followed by a median band of two vanishing-point tiles bisected by horizontal white lines. At the center lies a single approaching-point tile of a different phase with no bisecting line. This arrangement reveals the self-similar, fractal-like operation of the hierarchical matrix within the surface itself, demonstrating how alternating approaching- and vanishing-point phases propagate through successive scales to produce a coherent, wave-based relational structure.

The union of left and right waves into a polar continuum resembles certain physiological processes in the visual system, where the retinas transmit crossed and uncrossed phase signals that are reassembled in the visual cortex to support depth perception (16). In the GCC framework, alternating crossed and uncrossed surfaces similarly produce a continuous, balanced structure that enables perception of either approaching-point or vanishing-point perspectives.

This crossed/uncrossed strategy supports two key functions: enhancing depth perception through a visual polar continuum and helping identify the most hierarchical (and thus most parsimonious) geometric explanation among alternatives.

Because the GCC provides a coded template that each surface pattern must follow for accurate reproduction, the internal representation formed in the mind remains closely aligned with the external geometric structure it interprets. This alignment suggests that the framework may offer a unifying geometric principle linking discrete digital states with continuous wave-like perception and topographic surface behavior. This process may explain how an internal model of the external world is formed and maintained (17).

Thus, the mind internalizes not the objects themselves, but the connections between them. For example, at the atomic scale, one sublattice links directly to another along the Y-axis through warm and cool pixel shading (Fig.1). The topographic lattice model encodes the yin-yang relationship. The yin-yang relationship creates a balanced and harmonious whole, illustrating how contrasting forces such as light and dark, male and female, or active and passive, complement each other to maintain equilibrium (13,18)The GCC promotes balance by enabling inverted synchronization across lattices and waves, creating a unified structure.

## Conclusion

This study demonstrates a novel geometric complementary code (GCC) that bridges the discrete realm of pixel-based digitization with the continuous dynamics of wave mechanics. By oscillating four shaded cubic pixels within a Cartesian grid, the GCC enables a seamless transition to an emergent face-centered cubic (FCC) lattice structure across scales, incorporating the polarities of the sagittal, transverse, and coronal dissection planes. Expansion to larger Cartesian grids reveals hierarchical wave behaviors in which synchronization of complementary pairs produces constructive interference patterns along a yin-yang-like continuum.

The framework analyzes the geometric relationships of rise and run as topographic values within a surface grid. These components generate a polarized macro-grid when surface patterns repeat cyclically, enabling dynamic shifts between approaching-point and vanishing-point perspectives. The key insight is that polarities from the three dissection planes can be consistently applied to both lattice space and Cartesian grid space, thereby establishing a coherent hierarchical pixel-based morphometry.

The GCC offers a unifying paradigm that links discrete digital states with continuous topographic wave behavior. This approach holds potential for more efficient rendering of complex structures in computational imaging, graphics, material science, and biomedical visualization. Space appears most coherently understood as a succession of surfaces, and the GCC serves as a cohesive geometric force capable of bridging seemingly fragmented domains.

Future empirical validation in physical systems will be essential to test the GCC’s predictive power. Limitations include potentially high computational demands for large grids and assumptions regarding specimen symmetry.

D’Arcy Thompson proposed that grid-based methods in morphometrics could reveal non-genetic forces shaping biological diversity. The present pixel-based morphometry via the GCC affirms geometry’s foundational role in geometric morphometrics. By systematically encoding and grouping variations on a geometric theme, this framework validates and extends Thompson’s vision. In the economy of hierarchical systems, the GCC and its transforming grid reveal a hidden cohesive force that unifies disparate elements into a single geometric whole.

## Data Availability

The graphical data and source files supporting the findings of this study are deposited in the Zenodo repository at https://doi.org/10.5281/zenodo.18914890. During the peer-review process, the dataset is available via the private link provided in the cover letter. Upon acceptance and publication, the dataset will be made openly available under a CC BY 4.0 license.

## Statement on AI Assistance

During the preparation of this work, the author used *Grok* and *Copil*ot to make words clearer and more concise. After using this tool, the author reviewed and edited the content as needed and takes full responsibility for the content of the publication.

## References

1. Mitteroecker P. Thirty years of geometric morphometrics: Achievements, challenges, and the ongoing quest for biological meaningfulness. American journal of biological anthropology 2022:181–210.

2. Slice DE. Geometric Morphometrics. Annual Review of Anthropology 2007 September;36.

3. MacLeod N. What you sample is what you get: ecomorphological variation in Trithemis (Odonata, Libellulidae) dragonfly wings reconsidered. BMC ecology and evolution. 2022;22(1):43.

4. Spani F. Unveiling Nature’s Architecture: Geometric Morphometrics as an Analytical Tool in Plant Biology. Plants. 2025;14(5):80810339014050808.

5. Olsen TB, García‐Martínez D, Villa C. Testing different 3D techniques using geometric morphometrics: Implications for cranial fluctuating asymmetry in humans. American Journal of Biological Anthropology 2022 November;180.

6. Messer D, Atchapero M, Jensen MB, Svendsen MS, Galatius A, Olsen MT, et al. Using virtual reality for anatomical landmark annotation in geometric morphometrics. PeerJ 2022 February;10.

7. Waltenberger L, Rebay‐Salisbury K, Mitteroecker P. Three‐dimensional surface scanning methods in osteology: A topographical and geometric morphometric comparison. Am J Phys Anthropol 2021;174(4):846–858.

8. Bardua C. A Practical Guide to Sliding and Surface Semilandmarks in Morphometric Analyses. Integrative organismal biology. 2019;1(1):obz016.

9. Watanabe A. How many landmarks are enough to characterize shape and size variation? PloS one 2018;13(6):e0198341.

10. Vitkovic N, Radovic L, Trajanovic M, Manic M. 3d point cloud model of human bio form created by the application of geometric morphometrics and method of anatomical features: human tibia example. Filomat 2019 January;33.

11. Brombin C, Salmaso L. A Brief Overview on Statistical Shape Analysis. 2013 January.

12. Ben-Jacob ELH. The artistry of nature. Nature 2001 Feb 22;409(6823):985–6.

13. Fritjof Capra. The Tao of Physics. fourth ed. Boston: Shambhala; 2000.

14. Bertamini M. Processing convexity and concavity along a 2-D contour: Figure-ground, structural shape, and attention. Psychonomic bulletin & review. 2013;20(2):191–207.

15. Halliday D, Resnik R editors. Principles of Physics. 12th ed. Singapore: John Wiley & Sons; 2023.

16. Howard I, Rogers B. Binocular Vision and Stereopsis. New York, Oxford: Oxford University Press * Clarendon Press; 1995.

17. Lehar S. The World in Your Head: A Gestalt View of the Mechanism of Conscious Experience. Mahwah, N.J.: Mahwah, N.J. : Lawrence Erlbaum Associates, Publishers; 2003.

18. Wen-Ran Zhang. YinYang Bipolar Relativity. Hershey, PA: Information Science Reference; 2011.

